# Divided attention for feature-based selection in visual cortex

**DOI:** 10.1101/2020.01.27.922047

**Authors:** James C. Moreland, Geoffrey M. Boynton

## Abstract

Feature-based attention can select relevant features such as colors or directions of motion from the visual field irrespective of the spatial position. In visual cortex not only do we see feature-specific attention affecting responses in neurons with receptive fields at an attended location, but that effect also spreads to neurons with receptive fields beyond the spatially relevant location. When only one feature is task relevant, the spread of activity across space can act to facilitate perception of behaviorally relevant stimuli. However, when multiple features are relevant, what is the effect on behavior and brain activity? We tested this question by having observers divide attention between two patches (left and right of fixation) of moving dot stimuli, each containing overlapping upward and downward motion fields. In one condition, subjects performed a task on motion fields moving in the same direction in both patches. In another condition attention was divided between opposite directions of motion. Replicating a previous behavioral study, we found that observers showed better performance when dividing attention to the same directions of motion than opposing directions of motion. We analyzed the BOLD responses while observers performed this task using an inverted encoding model approach that provides estimates of responses to each of the four component dot fields. We found larger responses to the attended components, replicating previous studies of spatial and feature-based attention. However, these effects were much larger when attention was divided to between the same directions of motion than to opposing directions. Our fMRI results in area hMT+ predict our behavioral results by extending the normalization model of attention to include a global feature-based attention component in that leads to suppressed responses to attended stimulus components when attention is directed to opposing directions of motion.

## Introduction

Feature-based attention can select relevant features such as colors or directions of motion from the visual field irrespective of the spatial position (e.g. Boynton 2009; Martinez-Trujillo and Treue 2002; McAdams and Maunsell 1999). In human visual cortex not only do we see feature-specific activity acting at the location of an attended stimulus, but that activity also spreads to stimuli beyond spatially relevant location (Saenz et al., 2002). This spread of feature-based attention across space can act to facilitate perception of all stimuli containing the relevant feature. However, when multiple features are relevant across multiple stimuli, what is the effect on behavior and brain activity?

In the present study, fMRI responses were measured in the visual cortex while subjects divided attention between two patches (left and right of fixation) of moving dot stimuli, each containing overlapping upward and downward motion fields. In one condition, subjects divided attention across the two patches and performed a task on two fields moving in the same direction. In a second condition, attention was divided across space to opposite directions of motion. We replicated previous behavioral results using similar stimulus conditions and found that performance is better when attention is divided to the same feature than when dividing attention to different features (Andersen et al., 2013; Saenz et al., 2003). These behavioral results suggest that it is not possible to enhance the response to two features independently at two locations.

We applied an inverted encoding approach to our fMRI results (IEM: Brouwer and Heeger 2009; Foster et al. 2017; Sprague and Serences 2013) to examine the effect of attention on the response to each of the four dot fields. This approach enables a quantification of the population response to each of the two directions of motion at each of the two spatial locations. Our results show an overall larger response to the two attended stimulus components, replicating previous studies of spatial and feature-based attention. However, consistent with our behavioral results, in area hMT+ these attentional effects were much larger when attention was directed to the same directions of motion than opposing directions. Our fMRI results in area hMT+ can be predicted by an extension of the normalization model of attention which incorporates a spread of feature-based attention across space and a normalization process that suppresses responses to opposing motion directions.

## Methods

Methods were preregistered with the open science framework prior to data collection and can be accessed here: https://osf.io/hjm75/

### Participants

Seven healthy subjects (including author JM) with normal or corrected to normal vision were recruited from undergraduates and graduate students at the University of Washington. All subjects provided written informed consent and received $20/hr compensation (other than the author JM) for their behavioral participation and $30/hr for fMRI participation. Subjects completed 1-2 (number determined by reaching a level of acceptable performance) behavioral sessions lasting one hour, and 2-3 fMRI sessions each lasting 1-1.5 hours.

### Stimuli and procedure

#### Basic stimuli

Each stimulus component was a circular aperture of dots placed to the left and right of fixation (location parameters: 3.5° radius, centered ±6° away horizontally and −2.75° vertically) moving either upward or downward (dot parameters: white 100% luminance; speed 5°/s; 0.25° diameter; limited lifetime of 12 frames; 100% direction coherence) (**Figure 1B**). The display background was set at 30% of max luminance. Displays could contain one (forward model scans) or all four (divided attention task) component fields of moving dots (**Figure 1C**). Because all the dots were the same color within the aperture, the only cue for segmentation was the motion direction.

**Figure 1.**
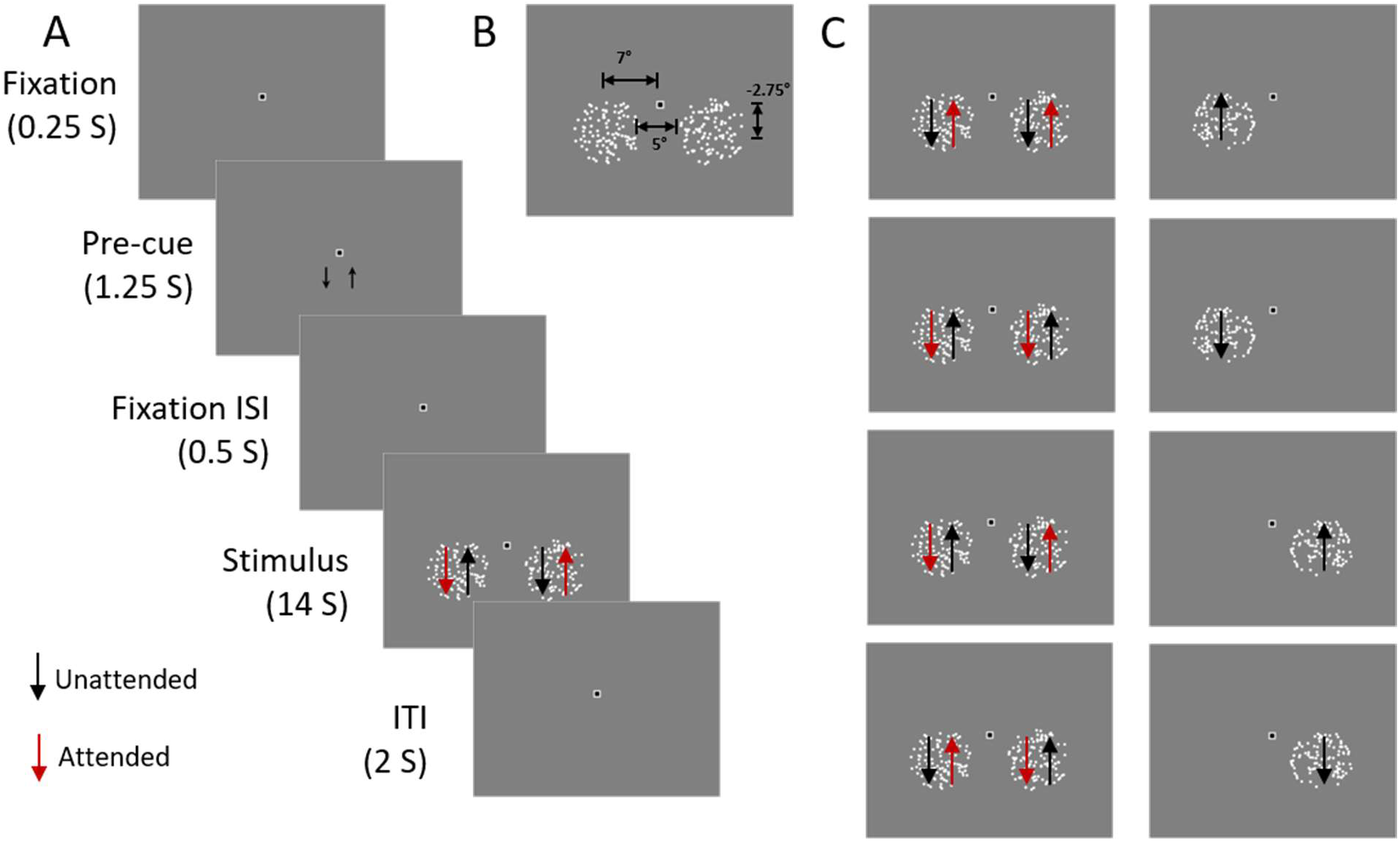
**A)** Schematic of an example trial in the divided feature attention task. The pre-cue could direct the observers to either field (up or down) on each side. There were no arrows on the stimulus display, these are included to indicate the directions of the four fields and the red indicates the relevant fields on this example trial. **B)** Stimulus size. **C)** Left column: Attention conditions. Red arrows indicates the relevant fields in each condition. Right column: Single field conditions used in forward model scans. There are no red arrows because attention was directed to a fixation task.

#### Divided attention task

Error! Reference source not found.**A** shows a schematic of the procedure applied during single trials for both fMRI and behavioral experiments. The stimulus display consisted of four fields of moving dots. Two overlapping in each aperture – one moving upwards and one moving downwards. The observers’ attention was directed to individual fields using arrows cuing a single field on each side (dual-task) as relevant to a luminance decrement detection task. The arrows could indicate fields that are moving in the same direction (*attend-same*), or in opposite directions (*attend-different*). There were four combinations of dot fields which could be attended per trial: up (on left)/up (on right), down/down, up/down, and down/up. Importantly, this task manipulated the attentional state of the observer while keeping the physical stimulus, eye position and spatial distribution of attention identical. Trials were completed as blocks of five trials of the same attention condition (e.g. left up, right up). Each attention condition block was completed once per scan for a total of 20 trials per scan.

Target events are defined as brief luminance decrements (from max screen luminance to background luminance of 30%, occurring in a Gaussian temporal window with SD = 200 ms) in the trial’s relevant fields. Distractor events are defined as luminance decrements of the dots in the trial’s irrelevant fields. On each 14 Sec stimulus presentation there were variable numbers of target and distracter events to ensure that the observer maintained attention throughout the trial and not only until they thought all target events had occurred. Events were randomly interleaved and spaced at least 750 ms apart with the first event occurring at least 500 ms after the dot onset and the last event ending 500 ms before the dot offset. It was made clear to the observers that the events were randomly timed and were independent across fields.

The subject began by foveating a fixation point on the center of the display and were required to maintain fixation on this point throughout the entire trial. At the start of each trial an arrow (1250 ms; 1° horizontal and 2.75° vertical displacement from fixation) on each side of the display indicated the relevant field directions for that trial. After a brief delay (500 ms), the moving dots stimulus was initiated and continued for 14 s. The observer’s task was to press a button when they detected a target event. Responses for a left field were made with the left hand, and responses to a right field were made with the right hand. The next trial will began after a blank inter-trial interval of 2 s. Trials were grouped in sets of 5 within an attention condition block. There was 10 s between each block. There was a total of 20 trials in each scan (394 s scans including a 4 s time before the first trial). The order of attention conditions was randomized within scans.

During training, feedback was given in the form of the fixation point turning green for correctly detecting (Hit) a luminance decrement in a relevant field and pressing the corresponding response key within 1 s. When an event was incorrectly detected or a response was made on the wrong side then the fixation point would turn red. In the behavioral only session, feedback was used until performance was at > 70% hits. Feedback was also used prior to beginning a scan session to remind observers of the task.

During practice in the laboratory, the stimuli were generated and displayed via CRT monitor with a 120 Hz refresh rate while eye position was recorded using an Eyelink II, 2.11 with 250Hz sampling (SR research, Ottawa, Ontario, Canada). During sessions conducted in the scanner, the stimuli were generated using a Dell XPS laptop and back-projected onto a fiberglass screen via an Epson Powerlite 7250 projector Stimuli for all experiments were created with Matlab software (MathWorks) and presented using the Psychophysics Toolbox (Brainard, 1997; Pelli, 1997).

#### FMRI data acquisition

fMRI data was acquired in a Phillips 3T Achieva MRI scanner at the Diagnostic Imaging Science Center at the University of Washington. Functional images were acquired using an echo planar sequence, with a 32-channel high-resolution head-coil with a repetition time of 2 s and echo time of 30 ms. Eighteen axial slices (80 × 80 matrix, 22-mm field of view, no gap) will be collected per volume (voxel size: 2.75 × 2.75 × 3.4 mm). Anatomical images were acquired using a standard T1-weighted gradient echo pulse sequence. From each subject we collected six standard retinotopic mapping scans, six forward model scans, and eight experimental scans across three scan dates.

#### Forward model scans

MRI sessions also included scans in which the observer viewed single fields of moving dots in isolation (left up, left down, right up, right down; Error! Reference source not found. **C**), which were used to estimate each voxel’s responses to each stimulus component separately for use in the inverse encoding procedure. Each dot field was presented 4 times for 10 s with 8 s blanks between field presentations and the order was randomized. Throughout these scans, observers performed a fixation dimming detection task. This encouraged the observer to hold a stable fixation, and maintain their attention at the fixation point.

#### Behavioral analysis

Observers made button presses when they detected luminance decrements in relevant fields. Responses were classified as hits, selection errors, misses, and false alarms. A hit was defined as a button press within one second of an event occurring in the relevant field on the same as the button. A selection error was defined as a button press made within one second of a distractor event, or a target event occurring on the opposite side to the button press. A miss was defined as no button press within one second of an event in a relevant field. A false alarm was defined as a button press when no event of either kind had occurred in the previous second.

Behavioral performance was assessed as rates of each of the four types of outcomes. We compared the rates of each kind of outcome across attention condition using an ANOVA with subject as a random effect.

#### fMRI analysis

We used standard phase-encoding retinotopic mapping procedures to define visual areas V1, V2, V3 and V4 (Engel, Glover, & Wandell, 1997). These areas were restricted to regions of interest (ROIs) using the forward model scans. This choice of localizer is not circular with the main analysis because voxels identified with the spatial localizer are not be biased by any attention condition to be examined in the experiment. MT+ was defined as a contiguous group of voxels lateral to the parietal-occipital sulcus and beyond the retinotopically organized visual areas that exhibited a large response to the moving dot stimulus versus blanks.

All preprocessing (anatomical-functional coregistration, slice-scan time correction, motion correction, and linear trend removal) was performed using BrainVoyager. We then imported preprocessed fMRI voxel time courses into Matlab for analysis with custom software. We applied GLMdenoise version 1.1 (Kay, Rokem, Winawer, Dougherty, & Wandell, 2013) to improve the signal-to-noise in our data using noise regressors derived from task-unrelated voxels. The results with and without GLMdenoise were compared using reliability measures orthogonal to the conditions of our experiments and GLMdenoise proved to be more reliable and so all further fMRI results used GLMdenoise voxel timecourses.

##### Eye movements

Our fMRI analysis relies on voxels representing the same spatial region throughout the experiment. This means that our observer’s ability to fixate well was paramount. Prior to scanning, observers underwent training to fixate and minimize eye movements. During the scans, we measured eye position which we used to assess observer’s fixation quality and discard trials in which their eye movements were excessive.

From the experimental scans, trials were excluded when more than one second out of the 14 second stimulus duration was spent outside of a 2° radius around fixation. One subjects was outside of the fixation ring for one second or greater on 62% of trials in one of their scan sessions, on this basis we did not have enough data in all of the attention conditions leading to the exclusion of this session from further analysis. For the remaining subjects, of the 120 trials each subject completed, 11% ± 5.7% were excluded from each subject. Exclusion of trials was implemented by regressing out that trial.

By comparison, in the single field scans, only 1.0% ± 0.6% of trials had periods greater than one second outside of fixation. During these scans the observer’s task was to detect luminance changes in the fixation point rather than in the fields because we wanted to keep attention off of the stimulus fields. It seems that observers were better able to maintain fixation under these conditions. This can explain why we find relatively high reliability in the single field voxel weights and more noisy weights in the attentional weights.

##### Inverted encoding procedure

We employed an inverse encoding procedure to estimate population responses, within each ROI, to each of the two spatial locations and directions of motion. Details are supplied in *Supplementary Material*. The forward encoding model assumes that each voxel contains subpopulations of direction selective neurons, and that there is an uneven balance in the overall direction and spatial selectivity of neurons contributing to a given voxel’s response (Kamitiani and Tong, Current Biology 2006). Using our fMRI responses we characterize a voxel’s overall direction sensitivity as a weighted sum of the four hypothetical channels (left side moving up, LU; left side moving down, LD; right side moving up, RU; right side moving down, RD). The first stage of this analysis uses fMRI data recorded while viewing displays of single fields of moving dots in isolation (LU, LD, RU, RD) to estimate weights of each voxel to motion direction and location. In the second stage, the fMRI data from the divided attention experiment was used with the estimated channel weights from above to estimate the response to each stimulus component, using linear regression under the assumption that the pattern of response our stimulus components is a linear combination of the responses measured to each stimulus component alone. The four regression weights serve as estimates of each channel’s response to each of the four stimulus components.

For each of our five regions of interest (ROIs: [V1, V2, V3, V4, and MT+]), in each of our subjects we obtained channel weights for each of the four attention conditions (attend LURU; attend LDRD; attend LURD; attend LDRU). After collapsing across scans+, this resulted in a four factor balanced data set: 7 (subject) × 5 (ROI) × 4 (weights) × 4 (attention condition) with ROI as a within-subject factor.

We hypothesized to find two results in a visual area containing motion information supporting the behavioral results. First, we expected that channel weights associated with the two attended stimulus components are larger than for the two unattended components – a main effect of attention, and (2) we expected a greater difference between attended and unattended channel weights for *attend-same* than the *attend-different* conditions – an interaction between attention and attention condition. We tested this by collapsing the 4 (channel weights) × 4 (attention condition) matrix of channel weights for each ROI and subject into 2 × 2 matrix of values (attend vs. unattended channel × *attend-same* vs *attend-different* attention condition). For example, the attended channel *attend-same* condition is the average of the left up and right up channels for the attend left up, right up condition, and the left down and right down channels for the attend left down, right down condition. The unattended channel *attend-different* condition is the average of the left down, right up channels for the attend left up, right down condition and the left up, right down channels for the attend left down, right up conditions. Thus, the 7×5×4×4 set of channel weights collapses to a 7 (subject) × 5 (ROI) × 2 (attended vs. unattended) × 2 (same vs. different) data set.

We ran a 2×2 ANOVA with ROI as a within-subject measure to test for (1) a main effect of attended vs. unattended, (2) a main effect of *attend-same* vs. *attend-different* condition and (3) an interaction between attended vs. unattended and *attend-same* vs. *attend-different*. Planned comparisons included simple effect ANOVAs for each ROI, again testing for the two main effects and interaction stated above.

## Results

### Behavioral results

Subjects performed a dual-task detecting luminance decrements on two fields. The fields were either moving the same direction or different directions. Luminance decrement events could either be correctly reported as present in a relevant field (hit), not reported in a relevant field (miss), reported when occurring in an irrelevant field (selection error - SE), or reported when not present in any field (false alarm - FA).

#### Behavior Only Session

**Figure 2** (top row) shows the results of the behavioral task during laboratory behavior sessions. There was no difference between *attend-same* and *attend-different* conditions for hit rate and false alarms (hit rate: F(1,6) = 0.48, p = .51, *η*^2^ = .97; false alarm rate: F(1,6) = 0.47, p = .52, *η*^2^ = .60). However, there was a significant difference between *attend-same* and *attend-different*

**Figure 2.**
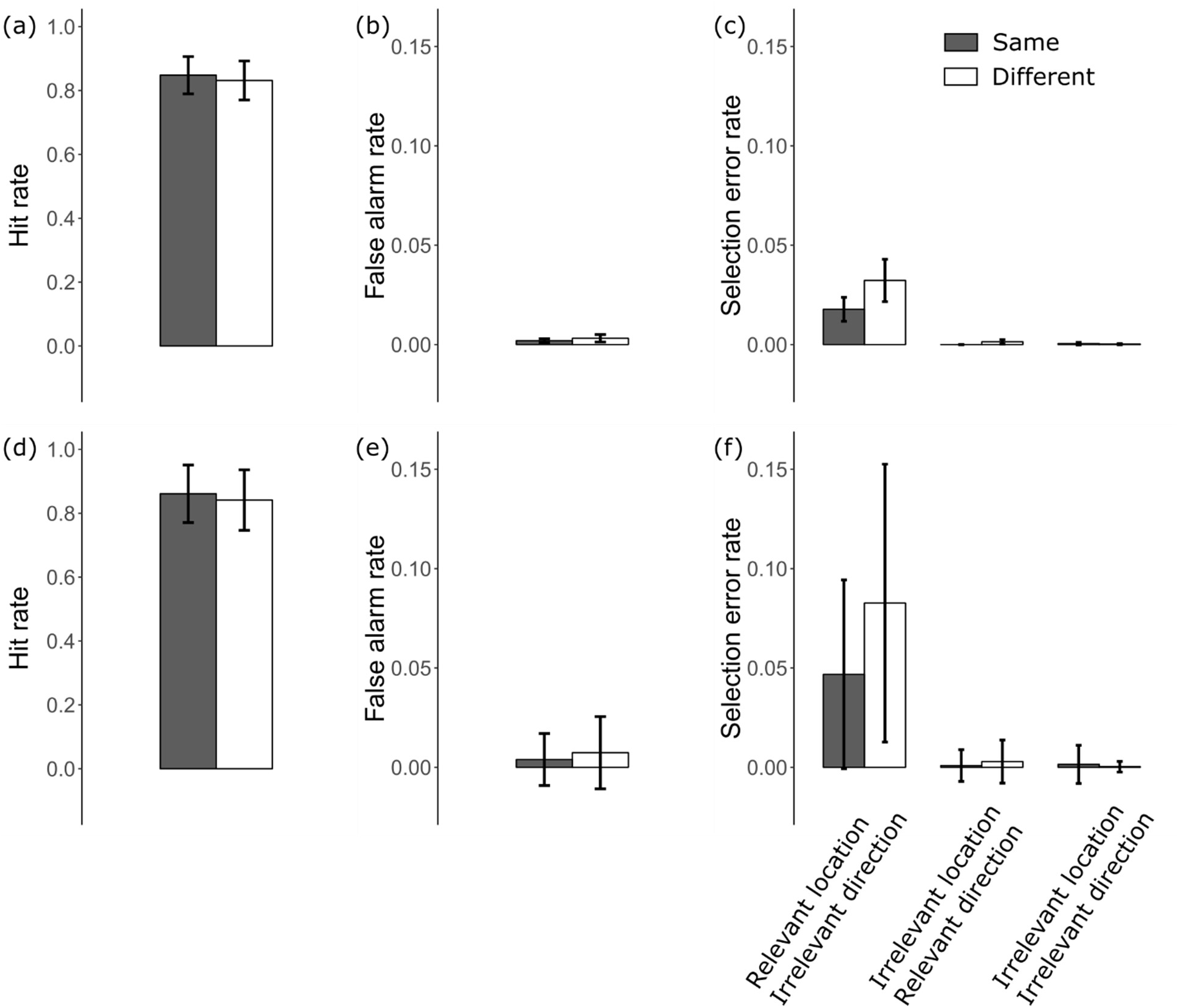
Results of behavioral task in behavioral only session (a-c) and in scanner sessions (d-f). Columns from left to right: Hits, False Alarms and Selection Errors. There are three kinds of selection errors. Responses made when an event occurred in the relevant location but in the irrelevant field, responses made when an event occurred at the irrelevant location in the relevant feature value, and responses made when an event occurred in an irrelevant location and irrelevant feature value. Error bars are standard error of the mean across subjects.

**Figure 3.**
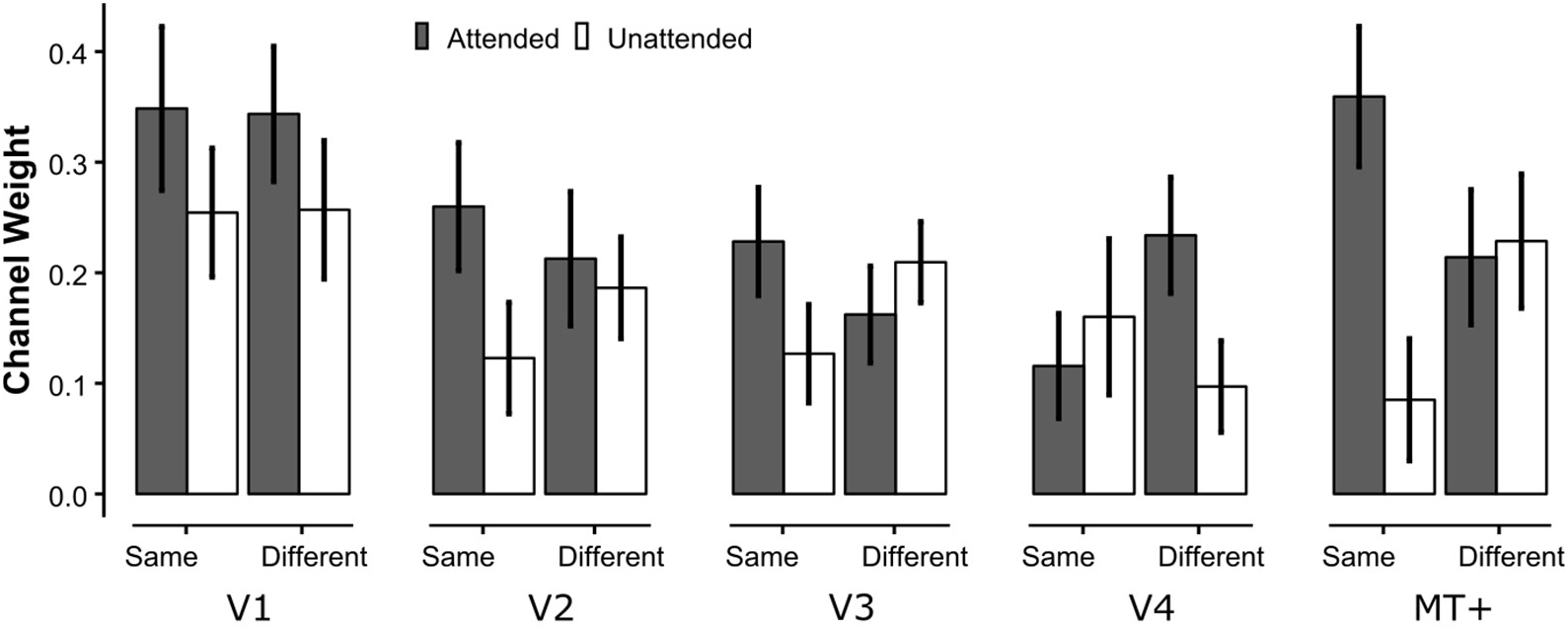
Channel weights for attended and unattended fields in the same and different conditions, for each visual area.

#### Behavior in the scanner

**Figure 2** (bottom row) shows the results of the behavioral task during scanning sessions. As in the laboratory, one-way ANOVAs with subject as a within factor found no difference between *attend-same* and *attend-different* conditions for hit rate (F(1,6) = 1.93, *p* = .21, *η*^2^= .98) or false alarms (F(1,6) = 1.25, *p* = .31, *η*^2^= .6). There was, however, a significant difference between *attend-same* and *attend-different* conditions in selection errors (F(1,6) = 7.37, *p* = .035, *η*^2^= .91).

### fMRI results

On each trial, two fields were attended (one on each side) and two fields were unattended (one on each side). There were also two kinds of same trials (LURU, LDRD) and two kinds of different trials (LURD, LDRU). In our analyses we collapse across these pairs of fields and pairs of attention trials to give us channel weights for same-attended, same-unattended, different-attended, and different-unattended.

**Figure 33** shows the channel weights from the inverse decoding method in each area showing the response to the attended and unattended stimulus component for each of the two attention conditions. We used a 2 (field: attended vs unattended) × 2 (attention condition: *attend-same* vs *attend-different*) ANOVA (ML) with ROI as a within subject factor with subject as a random effect. We found a main effect of field, no main effect of attention condition, but an interaction of field with attention condition (field: F(1, 102) = 15.0, *p* < .001), attention condition: F(1, 102) = 0.48, *p* = .49), interaction: F(1, 102) = 6.45, *p* = .01)).

This interaction between field and attentional condition is particularly strong in area hMT+, which shows a large attentional effect for the *attend-same* condition but no attentional effect for the *attend-different* condition. These channel responses in area hMT + are consistent with our behavioral results showing better performance for the attend same condition than the *attend-different* condition if we assume a signal-detection theory model in which performance on the motion task at a given spatial location is improved by both increased responses to the attended field, and decreased responses to the unattended field. Channel responses in area V1, show no interaction between field and attentional condition. These results would predict the same behavioral performance across the two conditions, inconsistent with our behavioral results.

### Normalization Model

We can predict the pattern of results in areas V1 and hMT+ using a modification of the normalization model of attention (Lee & Maunsell, 2009; Reynolds & Heeger, 2009), which is an extension of the general normalization mechanism suggested for cortical processing to account for effects of attention on neuronal responses (for review see Carandini & Heeger, 2012). **Figure 4** shows predicted responses when the excitatory drive is affected by attention in three ways. Model 1 (left column) is with no effect of attention on the excitatory drive (discs are uncolored). Model 2 (middle column) has spatial and feature-based attention operate independently, so only the attended two component responses are enhanced. We assume without loss of generality that in the *attend-same* condition, subjects are attending to upward motion on the left and the right. For the *attend-different* condition, subjects attend to up on the left and down on the right. Discs are saturated green for the two attended stimulus components and uncolored for unattended components for both conditions. Model 3 (right column) implements global feature-based attention for which the effect of feature-based attention spreads across space. For the *attend-same* condition, the excitatory drive is increased only for the two upward moving fields. For the *attend-different* condition, attention to upward motion on the left spreads to the upward component on the right, and similarly, attention to downward on the right spreads to modulate the excitatory response to downward on the left.

**Figure 4.**
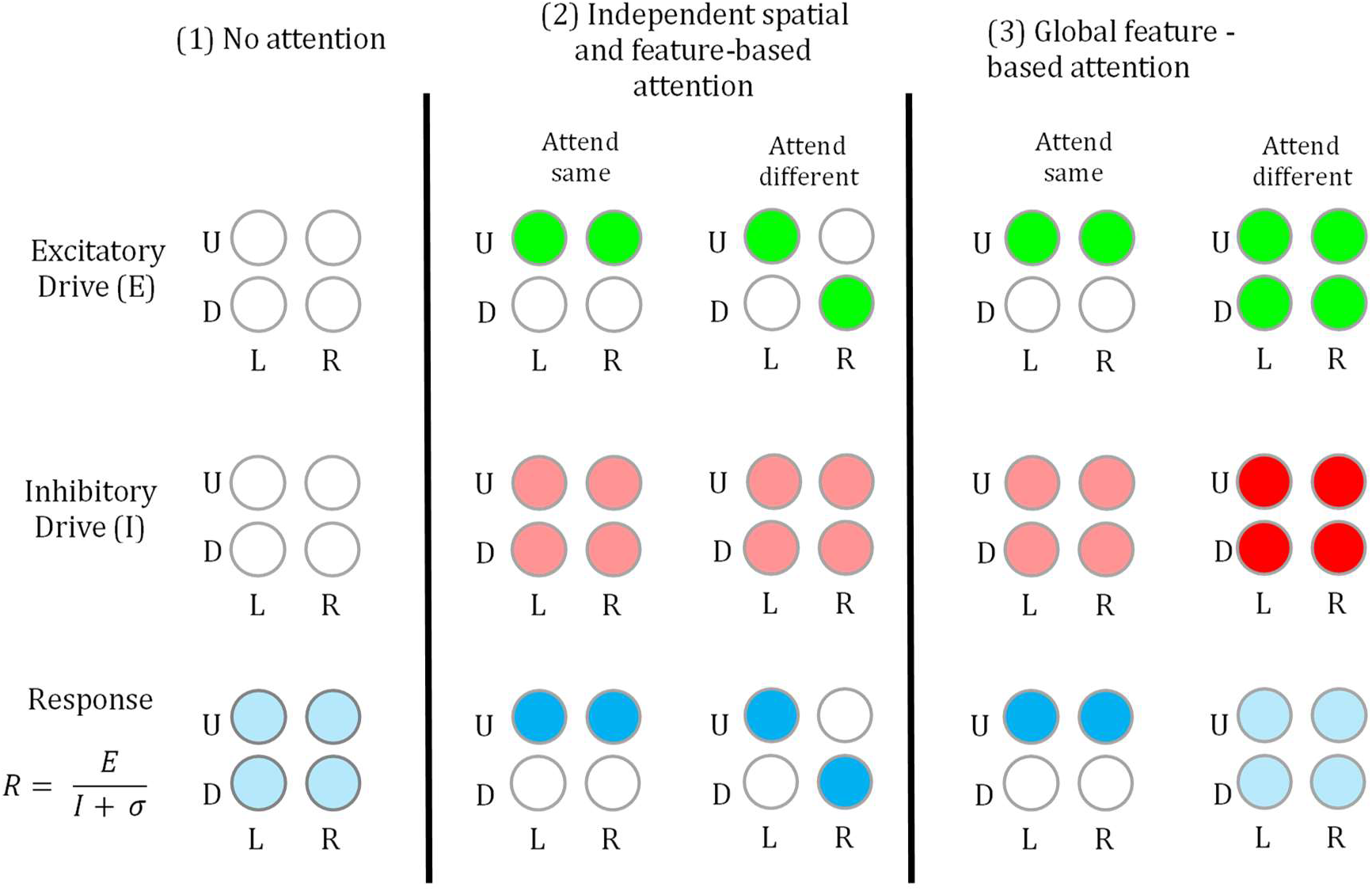
Illustration of how three implementations of how the normal model of attention can predict the effects of attention on neuronal responses. Discs represent responses to each of the four stimulus components (upward and downward motion in the left and right). The saturation of the color of each disc represents the strength of the response to that stimulus component along each stage of the model.

The next stage of the model computes the inhibitory drive, which is the excitatory drive summed across neurons tuned to all directions but with similar receptive field locations. In this illustration the inhibitory drive is the sum of responses across the two directions of motion for each spatial location. For model 1, there is weak excitatory drive for both directions and locations, so the inhibitory drive is also weak. For model 2, the inhibitory drive is intermediate, since it reflects the sum of the strong excitatory response to the attended component and weak excitatory response to the unattended component. Since spatial and feature-based attention are independent, the inhibitory drive is the same for both the *attend-same* and *attend-different* attention conditions. For Model 3, the inhibitory drive is also moderate for the *attend-same* condition. However, the inhibitory drive is strong for the ‘different’ condition because of the strong excitatory drive for both directions of motion.

The normalization model posits that the response in a population of neurons is the ratio of an excitatory drive, E, and inhibitory drive, I (plus a small constant, σ, to prevent infinite ratios). Spatial and feature-based attention only directly affects the excitatory drive – the rest of the processing in the model is the same, so that any effects of attention on the final responses are due to differences in excitatory drive.

The final stage essentially calculates the ratio of the excitatory drive and the inhibitory drive. Each of the three models predicts a distinct pattern of results. For model 1, the response is intermediate and equal for all four stimulus components, since this reflects the ratio of two weak responses. For model 2, if the effects of spatial and feature-based attention are independent, the response is large for the two attended stimulus components and weak for the two unattended stimulus components, reflecting the excitatory drive since the inhibitory drive does not vary across conditions. For model 3, with global feature-based attention, the effects of attention for the *attend-same* condition are the same as for model 2. However, the four responses for the *attend-different* condition are all intermediate and the same, due to the equal strength of the excitatory drive across stimulus components.

These model predictions can be sorted by whether the components are attended or unattended, and whether they came from the *attend-same* or *attend-different* condition to produce predicted channel weights that can be compared to our results. **Figure 5** shows predicted channel weights for the three models. Model 1, with no attention predicts no difference between same and different conditions and no effect of attention within these conditions. Model 2, with independent spatial and feature-based attention predicts that attended components have a larger response than unattended components but that there is no difference between *attend-same* and *attend-different* conditions. Model 3, with global feature-based attention shows an effect of attention in the *attend-same* condition but not in the *attend-different* condition.

**Figure 5.**
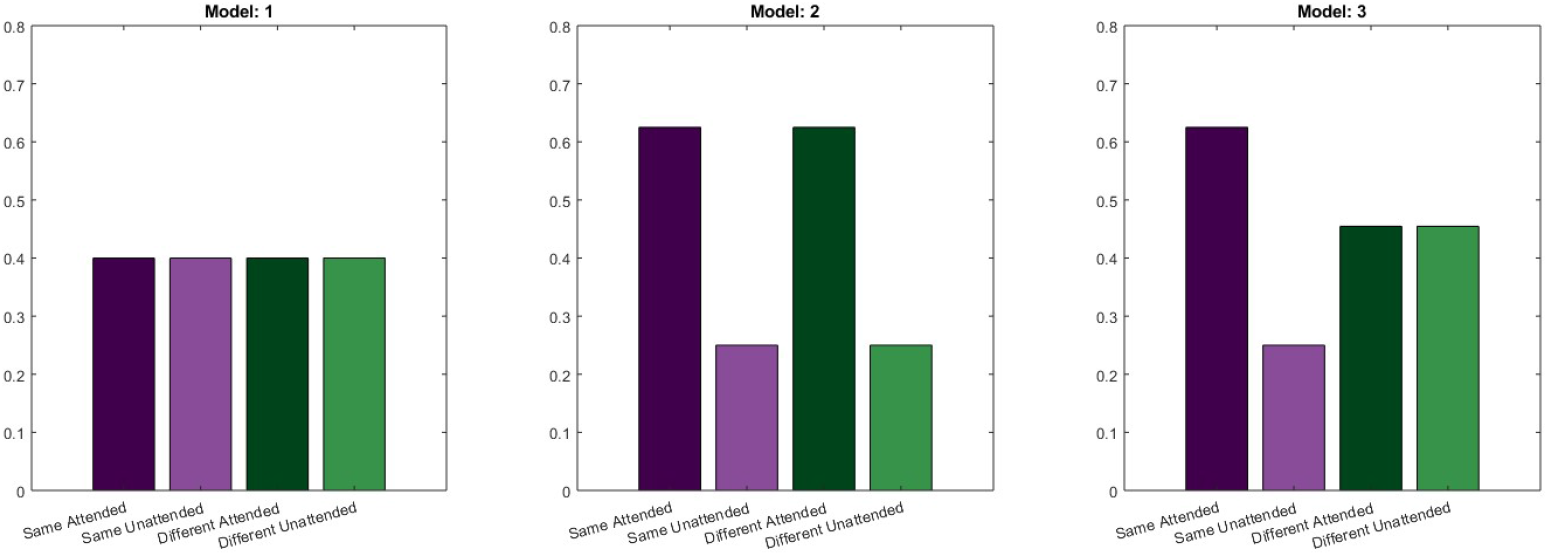
Predicted channel weights from the normalization model of attention for (A) no effect of attention, (B), independent spatial and feature-based attention, and (C), global feature-based attention.

Predicted responses from the three models can be sorted by attended and unattended components for the *attend-same* and *attend-different* conditions to predict channel weights for comparison to our fMRI results. Comparing the three model predictions to our fMRI results in **Figure 3**, it can be seen that our results in area V1 resemble Model 2, which has independent spatial and feature-based attention, and results in area hMT+ are consistent with model 3, which incorporates global feature-based attention.

Model 3 predictions, and area hMT+ results are consistent with our behavioral results, which supports the hypothesis that neuronal responses in area hMT + are used to ‘read-out’ the information needed to perform our divided attention task.

## Discussion

Can feature-based attention be constrained to a relevant location when divided across feature values? Behaviorally, we found that observers were able to detect events equally well when attending to different features at different locations as when attending to the same feature at the two locations. However, overall performance was compromised in the different feature condition by selection errors made within the same spatial location from the irrelevant stimulus field.

Our fMRI data supported a difference between same versus different feature conditions. Overall, feature-specific responses were greater for the relevant than the irrelevant fields. But there was a difference in the magnitude of this attention effect depending on whether the observer was attending to the same versus different field directions. When attending to the same direction, there was a larger response in the attended fields and a smaller response in the unattended field resulting in a larger difference between attended and unattended fields. These overall effects were driven by areas V1, V3 and MT+, which were the only areas to show the same effects when analyzed separately.

These divided attention effects have also been observed in the brain using EEG. Steady-state visually evoked potentials measured the responses to features at different locations and found enhanced activity for the when the same feature value was relevant at two locations relative to when conflicting feature values were relevant (Andersen et al., 2013; Andersen, Muller, & Hillyard, 2015; Forschack, Andersen, & Muller, 2017). This effect was greatly reduced, or eliminated, when the irrelevant features did not share feature values with relevant stimuli. These results suggest that there is feature-specific enhancement across the visual field but reduced responses to irrelevant features are specific to distractors and targets sharing feature values.

We see that there is only a significant difference for selection errors within the same spatial location between same and different conditions. This source of performance error was not clear in previous work where either percent correct from discrete trials (Saenz et al., 2003), or aggregated selection errors and false alarms (Andersen et al., 2013) were reported. This type of error is consistent with evidence that behavioral performance is only impaired when the distractors at each location match the relevant feature at another location (Andersen et al., 2013; Lo & Holcombe, 2014; Saenz et al., 2003). It seems that for selection of two features, the challenge is suppressing the attended feature at irrelevant locations, and failures result in selection errors.

Previous neuroimaging results have suggested the spread of feature-based attention beyond the relevant spatial location by showing feature related responses in regions outside of the physical stimulus (Saenz et al., 2002; Serences & Boynton, 2007a). Consistent with this, we find greater responses in the irrelevant fields when the attended direction is different on each side. This suggests that feature-based attention acts across the visual field which has the consequence of increasing responses to irrelevant fields sharing the same feature value.

An implication of this pattern of results is that even when it is not behaviorally advantageous to constrain feature-based attention to a local region this does not occur. Others have asked the question of whether global feature-based attention is obligatory. Some have suggested that the effects of feature-based attention depend on the task. Painter et al. (2014) showed using SSVEPs that there is only global feature-based attention gain when doing a conjunction visual search but not when doing a unique feature search. Andersen et al. (2015) instead find that feature-based attention occurs even when it is not advantageous for the behavioral task, and suggest that the discrepancy is due to the choice of baseline in the previous work. Our results agree that feature-based attention is not restricted even when it is to the detriment of performance.

The spread of feature-specific activity across space has often been described as a top-down gain control mechanism (Hayden & Gallant, 2009; Serences & Boynton, 2007a) which appears as a baseline increase in neural activity. The spread across hemispheres in our experiment could be due either to a global top-down mechanism, or a local top-down mechanism with hard-wired connections between similarly tuned neurons in each hemisphere. The experiment here cannot distinguish between these two possibilities but it is clear that our results cannot be fully explained with only a feature gain mechanism. Instead, we propose a simple extension to the normalization model of attention (Lee & Maunsell, 2009; Reynolds & Heeger, 2009). In addition to the spatial effect of attention, an effect of feature attention can be introduced by implementing increased activity across the visual field for relevant feature values. This method can predict the pattern of results we find in V3 and MT+ without having to add further parameters for capacity limits for dividing attention across feature values. Our results are not consistent with the pattern of results predicted by a normalization model with no effect of attention, or only spatial attention.

Reduction in responses during feature-based attention tasks with competing stimuli has be observed previously (Chapman & Störmer, 2018; Forschack et al., 2017; Müller, Gundlach, Forschack, & Brummerloh, 2018). Converse to global feature-based enhancement, feature-based suppression is thought to act locally to the relevant stimuli. For example, Forshcack et al. (2017) used SSVEP to determine the effect of feature-based attention on irrelevant stimuli in the periphery versus at the location of the relevant stimulus. They found that effects of feature enhancement could be detected in peripheral features matching the relevant feature value but there was no evidence of reduced responses in the periphery for the irrelevant feature value. We are not able to distinguish between local and global reduction in feature-specific responses, as the predicted results in our experiment would be the same. However, in MT+ it is likely that some voxels included neurons with receptive fields large enough to encompass both aperture locations. This would rule out a hypothesis of spatially isolated feature-based selection at least in area MT+.

Some studies have found evidence of feature-based attention in visual areas that we did not, such as V4 (Motter, 1994; Bichot, Rossi & Desimone, 2005; Buracas & Albright, 2009). Instead, we find an effect of feature-based attention in V3 where others did not (Serences et al., 2009). This is perhaps due to the attribute used in each study. It seems that the areas found to be sensitive to effects of feature-based attention are related to the attribute relevant in the task. V4 is more sensitive to color and shape, where V3 and MT are more sensitive to motion.

It should be noted that the observed effects of feature-based attention might not be due to modulations on neurons at early stages of processing (e.g. Liu et al 2007; Martinez-Trujillo & Treue, 2004; Maunsell & True, 2006) but from selective mechanisms further on in processing (e.g. Baldassi & Verghese, 2002; Palmer, 2005; Palmer, Verghese & Pavel, 2000; Dosher & Lu, 1999). This might reframe what appears to be enhancement and suppression as upweighting or downweighting of neural populations. It is possible that we are seeing these effects in early visual areas and are not able to distinguish between bottom-up and top-down because of the temporal resolution of fMRI.

### Concluding remarks

The present results show the effect of dividing attention across features on behavior and on responses in visual cortex. Our behavioral results show that it is more difficult to divide spatial attention to two different directions of motion than to the same direction of motion. Our fMRI results in area hMT+ are consistent with these behavioral results by showing a larger effect of attention when dividing attention to the same direction of motion than to different directions of motion. Our fMRI results can be predicted by a simple extension of the normalization model of attention, which incorporates a global spread of feature-based attention across space, followed by normalization process that pools responses to all directions of motion.

